# A cortical route for face-like pattern processing in human newborns

**DOI:** 10.1101/414284

**Authors:** Marco Buiatti, Elisa Di Giorgio, Manuela Piazza, Carlo Polloni, Giuseppe Menna, Fabrizio Taddei, Ermanno Baldo, Giorgio Vallortigara

## Abstract

Humans are endowed with an exceptional ability for detecting faces, a competence that in adults is supported by a set of face-specific cortical patches. Human newborns already shortly after birth preferentially orient to faces even when they are presented in the form of highly schematic geometrical patterns, over perceptually equivalent non-face-like stimuli. The neural substrates underlying this early preference are still largely unexplored. Is the adult face-specific cortical circuit already active at birth, or does its specialization develop slowly as a function of experience and/or maturation? We measured EEG responses in 1-4 days old awake, attentive human newborns to schematic face-like patterns and non-face-like control stimuli, visually presented with a slow oscillatory “peekaboo” dynamics (0.8 Hz) in a frequency-tagging design. Despite the limited duration of newborns’ attention, reliable frequency-tagged responses could be estimated for each stimulus from the peak of the EEG power spectrum at the stimulation frequency. Upright face-like stimuli elicited a significantly stronger frequency-tagged response than inverted face-like controls in a large set of electrodes. Source reconstruction of the underlying cortical activity revealed the recruitment of a partially right-lateralized network comprising lateral occipito-temporal and medial parietal areas largely overlapping with the adult face-processing circuit. This result suggests that the cortical route specialized in face processing is already functional at birth.

**Significance statement:** Newborns show a remarkable ability to detect faces even minutes after birth, an ecologically fundamental skill that is instrumental for interacting with their conspecifics. What are the neural bases of this expertise? Using EEG and a slow oscillatory visual stimulation, we identified a reliable response specific to face-like patterns in newborns, which underlying cortical sources largely overlap with the adult face-specific cortical circuit. This suggests that the development of face perception in infants might rely on an early cortical route specialized in face processing already shortly after birth.

## Introduction

As a highly social species, humans display a set of exceptional key competences for social interactions, that include the ability to detect, recognize and memorize faces, and to associate them with emotions and intentions (1). In the adult brain, face processing skills are coupled with a relatively highly face-specific set of cortical patches mainly localized in the right ventro-lateral occipito-temporal cortex (2, 3), but also extending to parietal, frontal and subcortical areas (4). Among those, the occipito-temporal face patches appear arranged in the same stereotypical pattern in humans, macaque monkeys (5) and even marmosets (6), suggesting a phylogenetic continuity in the primates’ neural systems underlying face processing.

Ontogenetically, a behavioral bias for faces is detected very early: human newborns within less than an hour from birth show a behavioral preference for canonically oriented faces, even when they are presented in the form of highly schematic geometrical patterns (two squares on top of one square, symmetrically inserted in an oval contour) over other kinds of visually controlled non-face-like stimuli (e.g., geometric patterns which configuration is incompatible with that of a face) (7–9). This early preference, observed both for schematic face-like configurations and real faces (10, 11), might already be present during the third semester of pregnancy (12) and it is also shared with other animal species like chicks and macaque monkeys (13–15). Preferential orientation to faces might be instrumental to increase newborns’ visual exposure to faces compared to other visual categories (16), providing the basis for rapidly developing specific face-processing skills.

What are the neural bases of this early bias for faces in the human baby brain? Is there a universally shared neural system that newborns deploy when processing faces versus other kinds of stimuli? The earliest evidence available to date for an early neural response to faces comes from EEG and fMRI/PET studies in infants of already 2-4 months. The EEG studies compared the Event-Related Potentials (ERPs) evoked by canonically oriented faces *vs.* inverted faces (17, 18) or *vs.* noise images with equivalent low-level visual properties (19), or contrasted the response to novel *vs*. familiar faces (20); in all cases, faces elicit a higher amplitude of the N290 and/or P400 ERP waves at occipital-temporal electrodes. A recent study in 4-6 months old infants using a novel EEG frequency-tagging paradigm (see more below), alternative to ERPs, confirms these results by showing a clear response to faces (compared with objects/scenes) in right lateral occipito-temporal electrodes (21). Although none of these studies attempted to reconstruct the anatomical sources of the EEG effects, their results are broadly compatible with the occipito-temporal neural generators of specific face processing ERP signals seen in adults (22). The rare fMRI/PET infant studies investigating the neural response to faces also confirm these results in 2 to 6 months-old infants (23, 24). The results are suggestive of an early cortical proto-architecture that preferentially engages when stimulated with faces. However, given the fast development of the visual system during the first three months (25), it remains an open question whether and to what extent the same occipito-temporal circuit involved in face processing in infants and adults is already active at birth, where the experience with faces is still extremely limited, or whether such specialization emerges only later as a function of experience and/or maturation.

Here, we aim to bridge this gap by investigating the electrophysiological correlates of processing face-like stimuli in awake, attentive human newborns of less than 96h after birth. We presented newborns with schematic and canonically oriented face-like stimuli (“upright face”) and, as controls, with an inverted version of the same stimuli (“inverted face”) (8, 9). As an additional control, we also presented “scrambled faces” organized in a non-face-like “top-heavy” fashion (more elements in the upper part than in the lower part of the oval) to investigate a previously proposed hypothesis that the preference for upright faces at birth may be mainly determined by a general preference for stimuli which geometrical organization is “top-heavy” versus “bottom heavy” (26).

In order to comply with the extremely short duration of focused attention in newborns (27), we took advantage of a frequency-tagging paradigm, a design that “tags” the neural populations coding for a given stimulus by presenting that stimulus periodically at a specific (‘‘tag”) temporal frequency and measuring the neural response in the form of a sharp peak in the EEG power spectrum at the same frequency (28). Since both the EEG ongoing activity and EEG artifacts are broad-band in frequency, the stimulus-related response in the frequency domain is easily discriminated from the stimulus-unrelated activity with relatively light artifact rejection, yielding a much higher SNR than the one obtained with ERPs. Oscillating visual stimulation based on the same principle has been widely used in the pioneer work on low-level visual function in newborns (e.g. (29, 30)).

We used a high-density (125 electrodes) EEG system (Electrical Geodesic, Inc, Eugene, OR) to record EEG activity in 1-4 days old healthy human newborns while presenting them with streams of schematic upright, inverted and scrambled faces (Fig. 1) presented periodically at a frequency of 0.8 Hz. Newborns’ stimulus-related brain responses were quantified from the peaks of the EEG power spectrum at the frequency of stimulus presentation. Cortical generators of the scalp-level effects were estimated with a source localization model based on newborn’s typical anatomical structure and electrical properties (see Methods).

**Figure 1:**
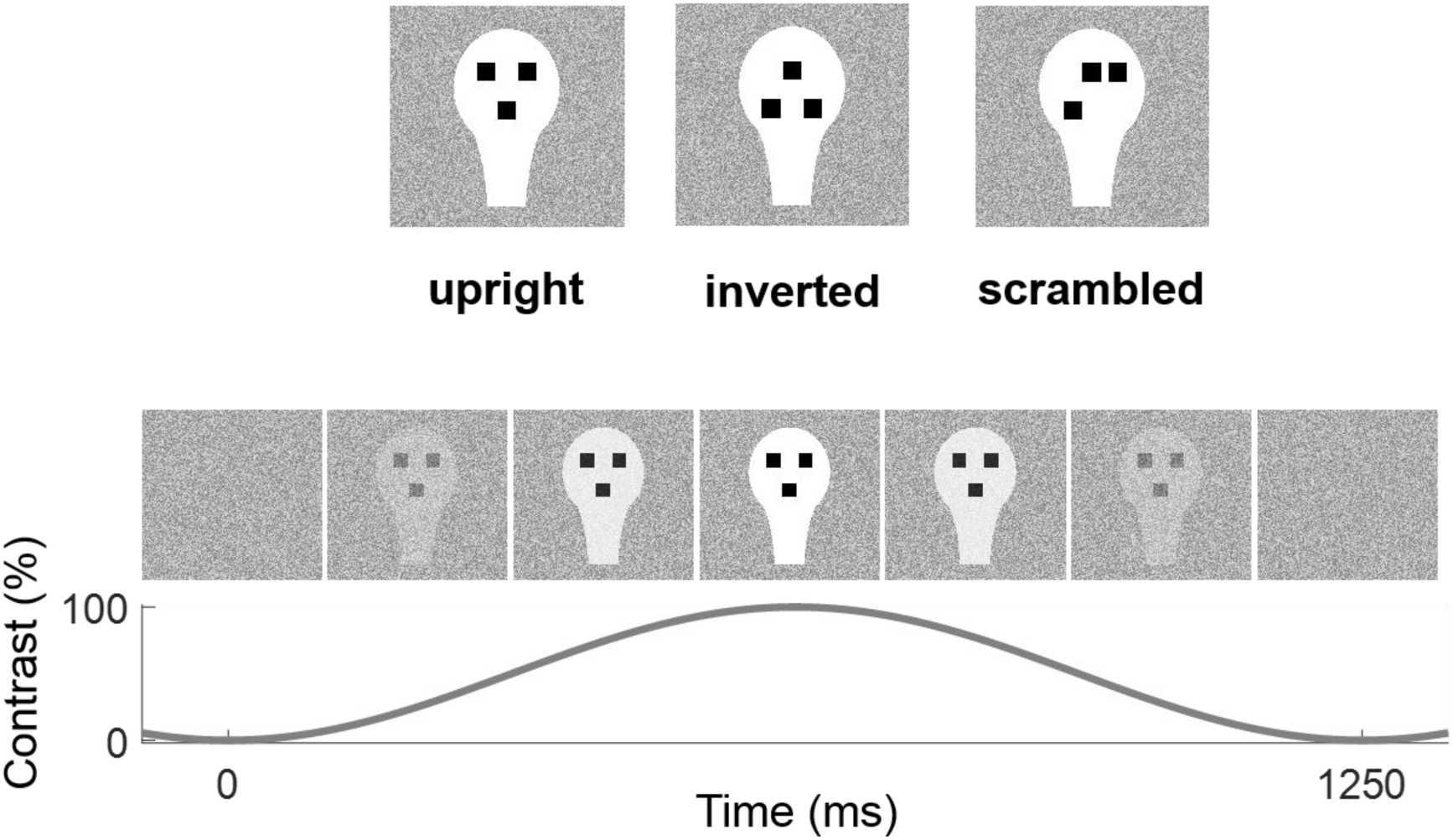
Visual stimulation. Top: Stimuli used (upright, inverted and scrambled face). Bottom: Illustration of one cycle of visual presentation with upright faces. Stimuli were presented dynamically with sinusoidal contrast modulation (0-100%) at a rate of 0.8 Hz (1 cycle = 1.25 s) overlapped onto a weakly contrasted dynamic background consisting of a flickering white noise image. Stimuli of the same type were presented continuously in blocks of 40 cycles (50 seconds) or until the subject stopped fixating.

## Results

Visual stimuli (upright, inverted and scrambled geometric representations of faces, see Fig. 1) were presented dynamically with sinusoidal contrast modulation (0-100%) in blocks of 50 s (or until the subject stopped fixating) at a rate of 0.8 Hz (1 cycle = 1.25 s), overlapped onto a weakly contrasted dynamic white noise background to minimize after-image effects. Data from the 10 subjects completing the protocol for all conditions were epoched on the basis of fixation intervals. After artifact rejection, the duration of clean EEG data per condition was on average 36.4 s, with no statistical difference among the three conditions (F(2,18)=0.28, *p*=0.68).

### All stimuli elicit a frequency-tagged EEG response

We first tested whether with such short data intervals we could reliably measure a significant oscillatory response at the frequency of stimulation. Given the steep 1/*f*-like profile of the power spectrum in the low-frequency range of the stimulation frequency in newborns (31), we estimated the stimulus-unrelated “background” power at the tag frequency by a power-law fit of the power spectrum at neighbouring frequency bins (±0.3 Hz). We then investigated the presence of a frequency-tagged response by testing whether and for which electrodes the power at the tag frequency was significantly higher than the estimated background power. Statistical testing for this as well as for all the following analyses was performed with a permutation-based non-parametric algorithm that tests the effects on the whole set of electrodes with no prior region-of-interest selection, where the issue of multiple comparison is overcome by directly assessing the statistical significance on spatial clusters of channels ((32), see Methods).

Results showed that taken together, the oscillating stimuli elicited at the tag frequency a significantly higher power than the estimated background power in a large set of posterior electrodes (*P*_*corr*_ < 0.003), and in a smaller frontal cluster (*P*_*corr*_ < 0.022) (Fig. 2A). Visualization of the power spectrum in the posterior cluster shows that, as expected, this effect is due to a high peak of power at the tag frequency emerging from a 1/*f*-like profile at neighbouring frequency bins (Fig. 2B).

**Figure 2:**
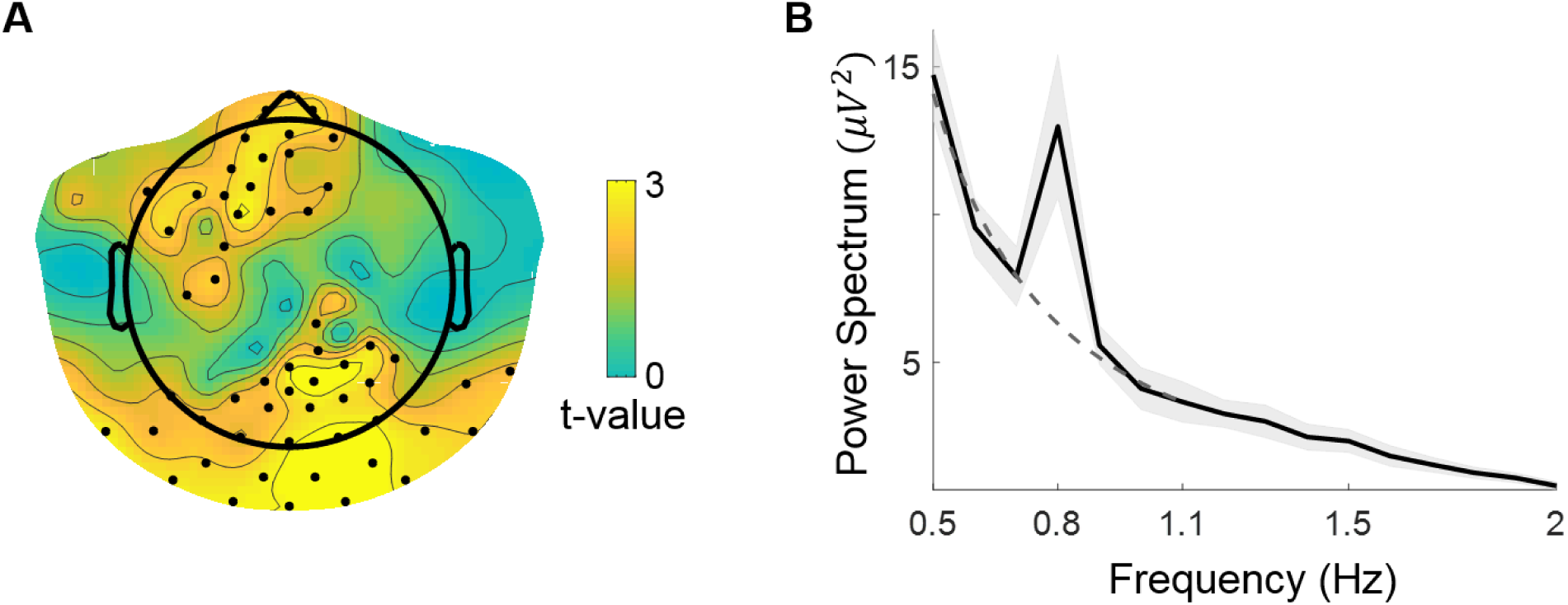
Frequency-tagged response, all conditions merged. (A) Statistical map (one-tailed *t*-test, corrected) of the difference between the power spectrum at the tag frequency (0.8 Hz) and the background power at the same frequency, estimated by a power-law fit of the power spectrum from the 6 neighboring frequency bins (±0.3 Hz). Electrodes belonging to a statistically significant cluster are marked with a black dot. Two clusters emerge: a posterior one (*P*<_*corr*_ 0.003) and a frontal one (*P*_*corr*_ < 0.022). (B) Power spectrum averaged over electrodes belonging to the posterior cluster (with *p*<0.01) (black line) ± s.e.m. across subjects (gray shadow): while the overall frequency profile is well described by a power-law (dashed dark-gray line, fitted in the interval 0.5-1.1 Hz), a peak neatly emerges at the tag frequency.

When taken separately, all stimuli elicited a significant peak at the tag frequency in a posterior cluster (upright: *P*_*corr*_<0.004; inverted: *P*_*corr*_ < 0.024; scrambled: *P*_*corr*_ < 0.020), while only upright stimuli gave rise to an additional peak in a frontal cluster (*P*_*corr*_ < 0.013) (Fig. S1).

### The electrophysiological signature of face-like pattern processing

Legitimated by the previous analyses, we quantified the Frequency-Tagged Response (FTR) to each kind of stimulus as the ratio between the amplitude of the power spectrum at the tag frequency and the background power at the same frequency, estimated by the power-law fit as above.

With that measure in hand, we moved to the direct investigation of the main question of our research: characterizing the electrophysiological signature of processing face-like patterns, by statistically comparing the FTR to face-like patterns first to inverted, and then to scrambled ones.

Compared to inverted faces, faces elicited a stronger FTR (Fig. 3B) in a wide posterior, slightly right-lateralized cluster (*P*_*corr*_ < 0.003) and in an anterior right-lateralized cluster (a weaker but significant effect; *P*_*corr*_ < 0.049) (Fig. 3A). Remarkably, the effect in the posterior cluster was consistently present in each single newborn (Fig. 4B). We denote hereafter as the face-like pattern response the subtraction between the response to upright faces and the one to inverted faces.

**Figure 3:**
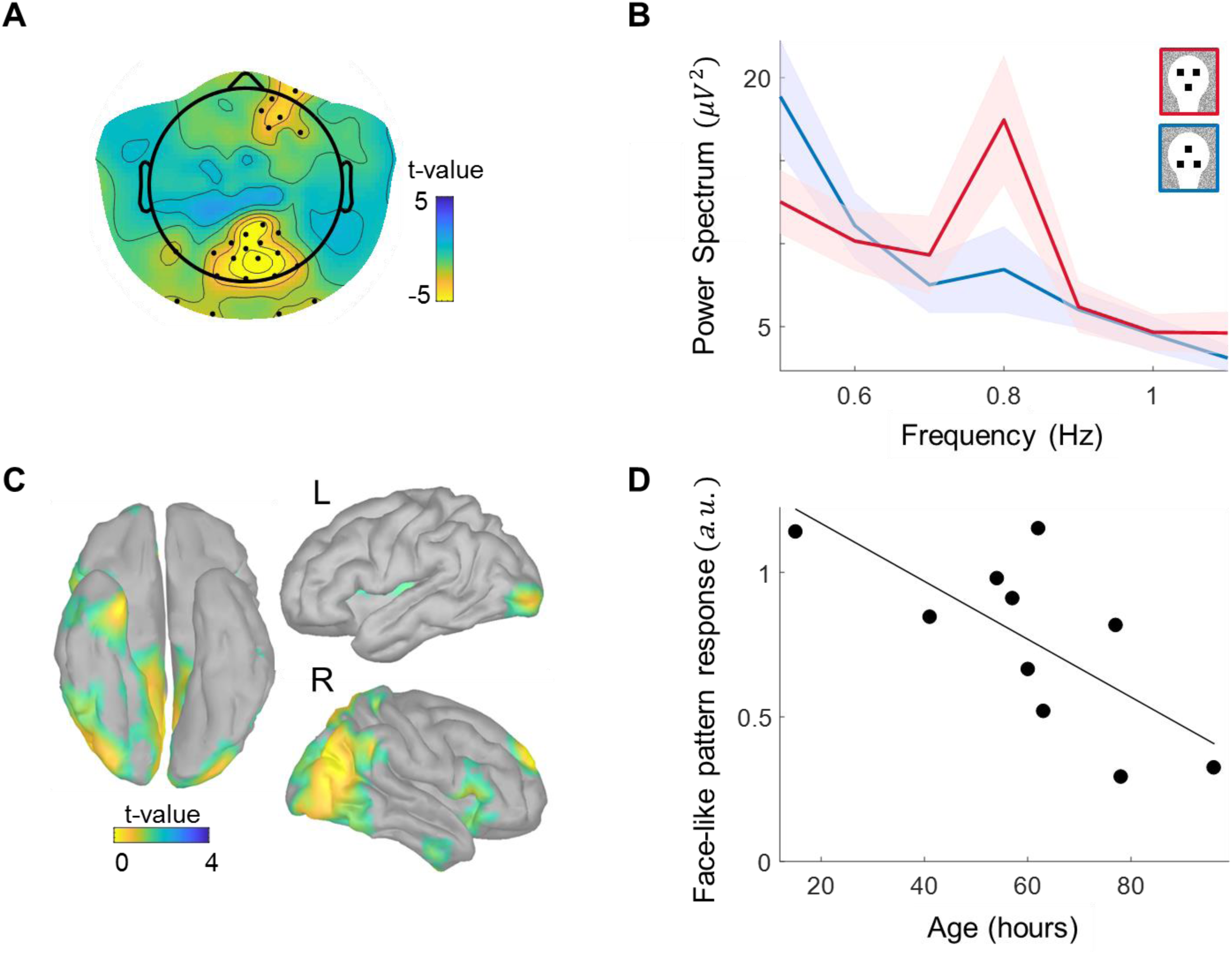
Comparison between upright *vs* inverted faces. (A) Statistical map (*t*-test, corrected) of the difference between the FTR to upright *vs* inverted faces. Electrodes belonging to a statistically significant cluster are marked with a black dot. Response to faces is significantly stronger in posterior (*P*_*corr*_ < 0.003) and right frontal (*P*_*corr*_ < 0.049) clusters of electrodes. (B) Power spectrum averaged over the posterior cluster (channels with *p*<0.01) for the two conditions (shaded contour indicates the s.e.m. across subjects): the tag frequency peak for upright faces is clearly higher than the one for inverted faces. (C) Statistical map of the comparison upright *vs* inverted faces at the source level (*p*<0.05, uncorrected), revealing a right-lateralized network that partly overlaps with the adult face processing network. (D) Inter-subject correlation between the face-like pattern response in the posterior cluster and the age from birth (R=0.71, *p*< 0.02).

**Figure 4:**
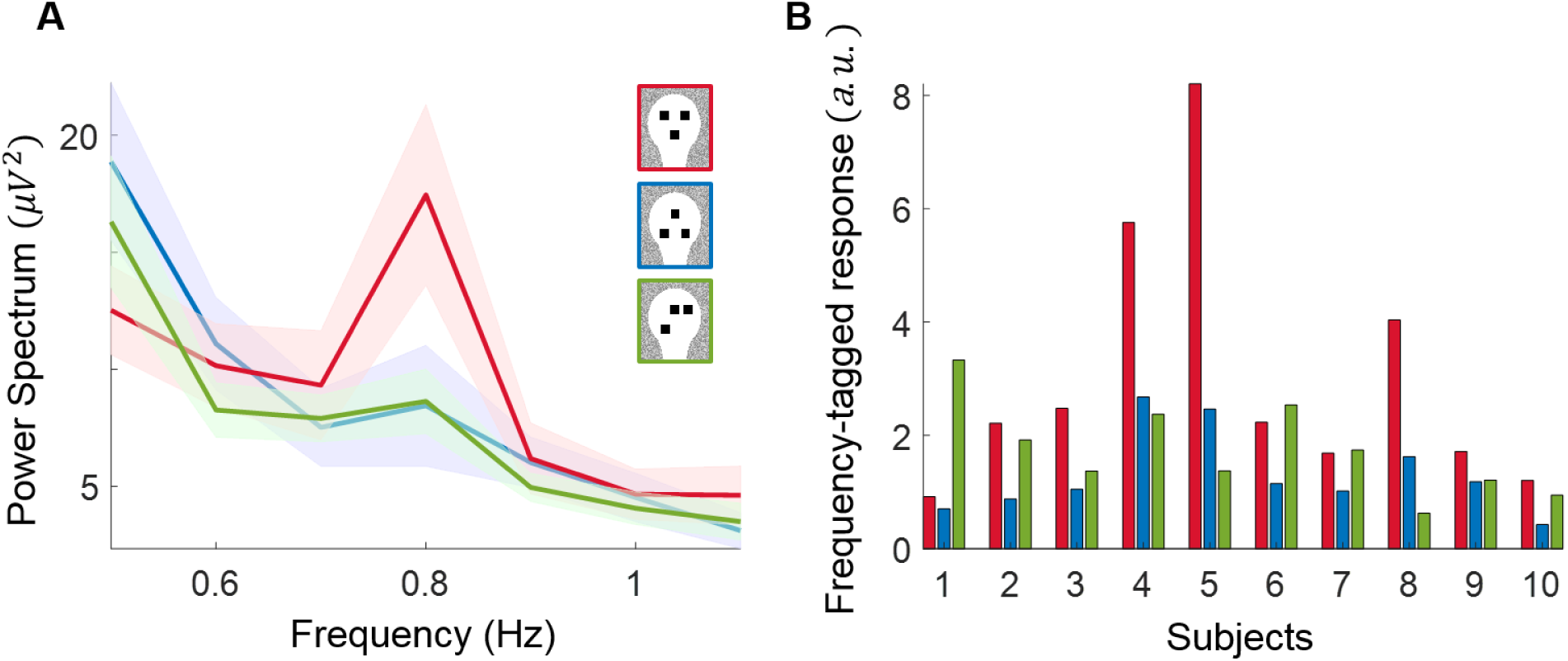
Response to scrambled faces is intermediate. (A) Power spectrum averaged over the posterior cluster associated to the face-like pattern effect (channels with *p*<0.01), for the three conditions (shaded contour indicates the s.e.m. across subjects): the average response to scrambled faces is more similar to the response to inverted faces. (B) Single-subject FTR in the same posterior cluster for the three conditions: while FTR(upright) > FTR(inverted) for each subject (reflecting the highly significant statistical difference), FTR to scrambled faces is closer to FTR to upright faces than to inverted in 4 out of 10 subjects, suggesting an intermediate response.

### The response to face-like patterns does not increase with age

In order to test the impact of age/exposure to faces on the face-like pattern response, we performed a correlation between age (in hours after birth) and the average face-like pattern response in the posterior and most significant cluster. Results showed a significant negative correlation (R=0.71, *p*< 0.02) (Fig. 3D).

### Estimated cortical sources of the response to face-like patterns

Capitalizing on the fact that since newborns have a higher skull conductivity than adults, a high spatial density sampling like ours (125-electrodes net) potentially captures significant spatial information (33), we estimated the cortical generators of the face-like pattern response identified at the sensor-level. We used an anatomical model morphed to newborns’ anatomy (34) to compute a detailed model of the infant head and cortical folds. We then used this forward model to reconstruct a plausible distribution of the cortical origins of our scalp recordings (see Methods).

The areas associated with the face-like pattern response at the source level (Fig. 3C) comprise a network that appears mostly lateralized to the right hemisphere, and that includes areas both along the occipito-temporal and the occipito-parietal stream: along the ventral stream activity emerges in bilateral occipital regions extending anteriorly to the right fusiform gyrus and the right ventral anterior temporal lobe, and superiorly towards the right posterior superior temporal sulcus. A strong activation was also seen in medial posterior regions including the right cuneus, precuneus, and part of lingual gyrus. Finally, some activation was observed in the right superior frontal gyrus.

### Response to scrambled faces is intermediate

We finally investigated the response to scrambled faces to test the hypothesis that they may yield the same pattern of response to faces due to their top-heavy configuration. However, contrary to the comparison faces *vs.* inverted face, the FTR to scrambled faces was not higher compared to inverted faces (no significant clusters, *p* > 0.05 for all uncorrected single-channel t-tests). On the other hand, FTR to scrambled faces was not significantly different from upright faces either (*P*_*corr*_ > 0.07). To further explore the nature of this intermediate response, we computed the FTR for scrambled faces in the posterior cluster associated with the face-like pattern response: while the average power spectrum is more similar to inverted faces than to faces (Fig. 4A), the response to scrambled faces is very variable across subjects (Fig. 4B), being in some subjects closer to faces (4 out of 10), and in others closer to inverted faces (6 out of 10).

## Discussion

### The mature cortical face network is present early in newborns

In this study we used a frequency tagging paradigm combined with high-density EEG to show that human newborns display a face-selective neural activation revealed by a higher response to face-like geometric patterns than to tightly controlled visual stimuli.

The estimated cortical sources of such response (Fig. 3C) extend along the occipito-temporal pathway, in areas closely resembling those found in most adult fMRI studies on face processing (occipital face area, posterior superior temporal sulcus face area, fusiform face area and anterior temporal face area; e.g. (2, 3)) as well as with intracranial EEG recordings using frequency-tagging in adults (35), and observed as early as in 2-4 months old infants (23, 24). These results suggest that the cortical face processing network is already laid down and functional in newborns.

In addition to this occipito-temporal right lateralized activity, we observe a medial activation centered in the precuneus, an effect similar to that described in face processing fMRI studies in adults when comparing familiar *vs.* novel faces (36). It thus might reflect the early-developed familiarity to a face-like pattern compared to non-face-like ones in newborns.

### Response to face-like patterns does not increase with age

While newborns spend most of their time sleeping, when they are awake the visual stimuli they are more frequently presented with are upright faces (16). One previous study from Farroni et al. indicated that the intensity of the near-infrared spectroscopy signal recorded in right occipito-temporal channels while newborns were viewing dynamic faces (39) indeed increased with age in infants from 24 to 120 hours after birth. This was taken as evidence that the cortical face-specific response requires frequent exposure to faces to develop. However, the specificity of the reported correlation is of difficult interpretation, as no control condition was used, leaving open the possibility that the increased activation to faces reflected a general maturation of the visual system. Indeed, our results are incompatible with the idea that the face-specific cortical response increases as a function of exposure to faces, as the correlation between age and the face-like pattern response is significantly negative (Fig. 3D).

In a speculative attempt to account for this surprising finding, we remark that we used highly simplified face-like geometrical patterns that for newborns act as “key” or “super-normal” stimuli (in ethological terms; see (37)), the sensitivity to which was previously shown to rapidly decrease already within the first month of life (Johnson 1991). One possible explanation is that while such key stimuli are optimally fit for the immature visual system of the newborn in the very first hours of life, experience with real world complex and variable faces may refine the face-like circuitry such that it rapidly gets more attuned to the real world features and gradually loses sensitivity to artificial face-like geometrical patterns. This fascinating but speculative possibility deserves further testing with a larger sample of newborns, e.g. by comparing the developmental trajectory of the cortical response to face-like patterns and real world faces.

### The role of cortical and subcortical structures

An influential theory proposes that newborn preferences for face-like stimuli may be mainly generated by a subcortical route involving the superior colliculus, amygdala and pulvinar (38). This theory, however, mainly relies on the assumption that the cortical visual route compared to the subcortical one is very immature in newborns, and on indirect behavioral evidence that the face preference phenomenon occurs only through the temporal visual route (39). One alternative possibility is that the processing of visual information proceeds in multiple waves of activation that involve – even the most rapid ones - both subcortical and cortical structures (40), an hypothesis supported by the dense connectivity of the structures of the subcortical visual route with multiple areas of the cortex (40), and by recent evidence from resting state fMRI studies that by term age the newborn cortex has reached a highly organized functional architecture (41) and thalamocortical connectivity (42), similar in many aspects to the adult ones.

Our study cannot provide evidence for or against an involvement of the subcortical route in face processing. In fact, because subcortical structures generate extremely weak electrical fields due to their closed-field geometry, and they are far from the scalp, they hardly produce measurable signals at the scalp level (43). We therefore believe that an impact of subcortical activity on our EEG results is unlikely and thus we did not include subcortical areas in our source reconstruction analysis.

On the other hand, our source reconstruction results support the hypothesis of a recruitment of a specific set of cortical structures in face-like processing at birth. Since this network widely overlaps with the adult face-processing circuit, we further speculate that one or more of these cortical areas might be already sensitive enough to face-like stimuli to generate the orientation preference to face-like patterns observed in newborns (7, 8). It is worth noting that this cortical recruitment is fully compatible with an early temporal subcortical route of the visual input (39), alternative to the relatively immature LGN/primary visual cortex pathway, as the pulvinar and amygdala are densely connected with (and massively influenced by) multiple cortical areas (40).

### Sensitivity of the EEG frequency-tagging paradigm over behavioral measures

Interestingly, while the early behavioural preference for upright face-like patterns compared to inverted ones is systematically observed in newborns by using preferential looking paradigms (where two different stimuli are concurrently shown on the screen, e.g. (9)), with single central presentations similar to the one used in the current stimulation paradigm, a behavioral preference for faces over non-face-like controls is typically not detected until 2 months of age (44). However, EEG responses to our centrally presented single stimuli did indicate a strong FTR difference across conditions, suggesting that in this case direct brain measures can be more sensitive compared to behavioral measures.

### Faces or top-heavy configurations?

Another result of the current experiment is that of an intermediate response of scrambled faces compared to faces and inverted faces. The fact that scrambled faces did not elicit a stronger FTR than inverted faces does not support the hypothesis that face preference reflects a preference for top-heavy configurations (26). In other words, the presence of a top-heavy configuration alone is not sufficient to systematically elicit a face-like neuronal response. On the other side, even if on average faces elicit a higher FTR compared to scrambled faces, the high inter-subject variability suggests that top-heavy stimuli may be sometimes categorized as a face.

### Future directions

Newborns spend most of their time sleeping, and during the rare periods in which they are calm and awake their visual attention typically lasts no more than 3-5 minutes (27). Here we show that the frequency-tagging paradigm provides a valid tool for measuring in newborns high SNR brain responses to multiple stimulus-specific conditions with very short stimulus presentation (around 40 s per condition), confirming results obtained with older infants (21, 45), and opening the way to investigate the neural substrates of other core perceptual/cognitive functions in this very special population. The high statistical significance of the face-like bias effect, reflected in its presence for each single subject, suggests that our experimental protocol, even in its shorter version limited to the presentation of upright and inverted faces, might be used as a biomarker to test the sensitivity to face-like patterns in populations at risk like ASD, as a complement to behavioural tests on visual social predispositions (46).

## Materials and Methods

The study was approved by the local ethical committee for clinical research (Comitato Etico per le Sperimentazioni Cliniche dell’Azienda Provinciale dei Servizi Sanitari della Provincia Autonoma di Trento, Resolution n. 1252|2015) and was performed in the maternity ward of the Ospedale Santa Maria del Carmine, Rovereto, Italy. Parents were informed about the content and goal of the study and gave their written informed consent.

### Subjects

Ten newborns (6 males; mean age 60 ± 22 hours, range 15 to 96 hours) were included. All were healthy (APGAR(1 min) ≥ 8, APGAR(5 min) = 10 for all subjects), born full term (gestation age: 39,7 ± 1,5 weeks), and of normal birthweights (average weight 3,41 ± 0,28 Kg).

44 additional newborns participated but they were excluded either because they did not complete the study (criteria: attend all three stimulus conditions for at least 20 s each) due to inattentiveness (18), falling asleep (16), crying (5), or because their data contained too many EEG artefacts (mainly due to movements or high electrode impedance) (5).

### Stimuli

Stimuli were presented using the Psychtoolbox 3.0.12 for Windows in Matlab R2014a (Natick, MA). Visual stimuli (Fig. 1, top panel) consisted of a white head-shaped form, 14.2 cm × 22 cm (25° × 38°) containing three black squares (2.5 cm × 2.5 cm, 4.4° × 4.4°), and differed only for the spatial configuration of the three squares. In the face stimulus (F), the squares were placed in the appropriate location for the eyes and the mouth to form an upright face-like pattern resembling a schematic human face; in the inverted face stimulus (IF), the spatial configuration of the squares was rotated by 180° to form an inverted face-like pattern; the scrambled face stimulus (SF) was obtained from the face stimulus by shifting the two upper squares on one side and the lower square on the opposite side of the head shape (sides were counterbalanced across subjects).

Stimuli were presented dynamically with sinusoidal contrast modulation (0-100%) at a rate of 0.8 Hz (1 cycle = 1.25 s) overlapped onto a weakly contrasted dynamic background (Fig. 1, bottom panel) consisting of a flickering white noise image (a rectangle, 45 cm x 33.8 cm, where the color of each pixel varies randomly between mid-gray (b/w intensity = 128/256) and grayish white (b/w intensity = 223/256) at a frequency of 3.75 Hz). We used sinusoidal contrast modulation instead of a squared on-off dynamics both to minimize non-linear effects in the brain frequency response (28) and to make the stimulation more pleasant to the babies (21). The slow presentation rate (0.8 Hz) was chosen to insure that newborns fully perceived and processed the stimuli within a sort of continuous “peekaboo” game.

### EEG recordings

EEG was recorded with a high-density (125 electrodes) Geodesic EEG system (GES400 EGI, USA) referenced to the vertex. Scalp voltages were amplified and digitized at 250 Hz.

### Experimental protocol

Newborns were tested in a calm, dimly illuminated space in the maternity ward, seated on the lap of a trained researcher in front of an 60 cm x 33.8 cm LCD screen (distance eyes-screen: about 30 cm) while wearing the EEG cap. Video recording from a hidden camera on the top of the screen insured on-line monitoring of the infant. The newborn’s parents, when present, were off the sight of the infant (separated by a curtain), and instructed to keep silent during the recordings.

Each trial started with a distracter consisting in a grey spiral looming towards the center of the screen on a reddish background. As soon as the newborn started to fixate the center of the screen, stimulation started with 1s of flickering background, followed by the periodic presentation of one of the three stimuli. Each condition was presented for 40 cycles (50 seconds) or until the subject stopped fixating and became bored or fussy. The trial ended with 1 s of flickering background followed by a blank screen. For each subject, the three conditions were presented in random order (counterbalanced across subjects). To maximize the EEG data statistics for all conditions, if the newborn kept her/his attention after each triplet of conditions, the same triplet was presented again, up to three times.

Fixation intervals were recomputed off-line by an experienced researcher (E.D.G.) who reviewed the video recordings blindly with respect to the experimental conditions. Newborns had an average fixation time of 43.4 s for condition. There was no statistical difference among fixation intervals in the three conditions (F(2,18)=0.24, *p*=0.74)).

### EEG data analysis

Data analysis was performed with the EEGLAB toolbox ((47), http://www.sccn.ucsd.edu/eeglab/), the Fieldtrip toolbox ((48), http://www.fieldtriptoolbox.org/), Brainstorm ((49), http://neuroimage.usc.edu/brainstorm/) and custom-made software based on MATLAB R2016b (Natick, MA).

### EEG pre-processing

EEG data were band-pass filtered between 0.1 and 40 Hz and segmented in blocks corresponding to fixation intervals, excluding intervals shorter than 10 s (i.e. eight stimulation cycles, the minimum duration required for the power spectrum analysis, see below). Bad channels were identified with the help of the TrimOutlier (https://sccn.ucsd.edu/wiki/TrimOutlier) toolbox by excluding channels that had a standard deviation higher than 150 μV or lower than 1 μV or that showed artefactual patterns at visual inspection (on average 4.9 per subject, min 0, max 14). Signals of bad channels were replaced with interpolated signals from neighbouring channels (standard spherical interpolation method in EEGLAB). Data segments containing amplitudes exceeding ±200 μV or containing paroxysmic artifacts after visual inspection were rejected. The resulting signals were mathematically referenced to the average of the 125 channels.

### Frequency-tagging analysis

In order to obtain a high frequency resolution of the power spectrum with one bin centered on the stimulation frequency (0.8 Hz), epoch length was set to exactly 8 stimulation cycles (10 s), resulting in a frequency resolution of 0.1 Hz. EEG data from each block were segmented in partially overlapping epochs of 10 s (overlap varied between 1/2 and 3/4 of epoch length to include all time points). For each electrode, the Fourier transform *F*(*f*) of each epoch was calculated using a fast Fourier transform (FFT) algorithm (MATLAB, Natick, MA). The power spectrum was calculated from these Fourier coefficients as the average over epochs of the single-epoch power spectrum: *PS*(*f*) = ⟨*F*(*f*) × *F* ^∗^(*f*)⟩_*ep*_. The Frequency-Tagged Response (FTR) at the tag frequency (0.8 Hz) was calculated as the ratio between the power spectrum at the tagged frequency and the value at 0.8 Hz of the power-law fit of the power spectrum estimated from the 6 neighboring frequency bins (+-0.3 Hz), where the power-law fit was computed by fitting a line to the logarithm of the power at the 6 neighboring frequency bins (Matlab function Polyfit). It is worth noting that, due to the steep 1/*f*-like power law of the power spectrum in newborns in the low frequency interval analyzed here (0.5-1.1 Hz) (31), the popular method to estimate the background power spectrum at the tag frequency by simply averaging over neighbouring frequency bins (50) overestimates the background power (and therefore underestimates the FTR) because the power spectrum is much steeper for lower than for higher frequency bins around the tag frequency.

### Statistical analysis

The frequency-tagging effect was evaluated, both for each condition and for all conditions merged, by comparing the logarithm of the power at the tag frequency with the logarithm of the background power estimated by the power-law fit described above.

Differences between the frequency-tagged responses across conditions were evaluated by comparing the logarithm of the relative FTRs.

In order to statistically evaluate the aforementioned effects we used the non-parametric cluster-based test (32) implemented in Fieldtrip (48). This method allows statistical testing with no need of a priori selection of spatial ROIs because it controls for multiple comparisons by clustering neighboring channel pairs that exhibit statistically significant effects (test used at each channel point: dependent-samples *t* statistics, threshold: *p*=0.05, one-sided for the frequency-tagging effect, two-sided for the differences between conditions) and using a permutation test to evaluate the statistical significance at the cluster level (Montecarlo method, 2000 permutations for each test). Results on statistically significant clusters are reported by specifying the polarity of the cluster (positive or negative) and its *p* value, indicated as *P*_*corr*_ to mark that it is “corrected” for multiple comparisons.

### Source reconstruction

As a head model, we used the one described in (34) (details therein). In brief, a realistic head model was generated from the anatomical magnetic resonance images (MRI) of a healthy full-term baby. The three-shell physical model included scalp, skull and intracranial surfaces downsampled to 2562 equidistant vertices each. Co-registration of the position of the EEG electrodes with the model was performed by using the co-registration of the same set of electrodes (EGI’s 125 electrodes set) with a slightly bigger (7-weeks-old) infant anatomy (51): we used Brainstorm tools to project such co-registered locations on our newborn head model. Since an accurate segmentation of the cortical gyri was not possible (due to contrast and resolution issues), a standard gyrated cortical surface (‘Colin 27’ in the Brainstorm software) was used as the source space in the model (52). Such cortical surface was rescaled into the infant brain size, smoothed to match with the cortical folding of a newborn (smoothing with a factor of 20%), positioned to the original cortical surface in the individual’s MRI and down-sampled to 8014 vertices. The forward model was computed by using the Symmetric Boundary Element Method implemented in the OpenMEEG software (53). Based on recent simulation and empirical studies on newborn head models, we set the following conductivity values: scalp 0.43 S/m, intracranial volume 1.79 S/m, and skull 0.2 S/m (33, 52, 54).

In order to obtain an estimate of the noise in the EEG signals that did not include any stimulus-related brain activity, we identified for each subject artifact-free data segments beginning 750 ms after the offset of each trial and ending at the onset of the distracter starting the following trial. Noise covariance was computed from these segments after application of the same pre-processing procedure used for the stimulus-related data.

To explore the anatomical sources of the statistically significant effects observed at the sensor level, FTR was estimated at the source level by performing the following steps: (a) For each subject, source-level time series were reconstructed from the segmented EEG data on the 8014 sources obtained from the wMNE reconstruction in Brainstorm; (b) Log(FTR) was estimated at the source level by using the same frequency-tagging analysis used at the sensor level; (c) Source log(FTR) values were spatially smoothed (10 mm).

For each contrast of interest, a paired *t*-test was run at each source location and the corresponding significant clusters (*p*<0.05 uncorrected) are reported on a template cortex smoothed at 50%. Importantly, the *t*-test at the source level is only used to properly describe the source distribution of the statistically significant effect established at the sensor level, and not for a second statistical test at the source level, therefore no correction for multiple comparison is required (55).

## Acknowledgements

We thank the staff of the Maternity Unit of the Rovereto Hospital S. Maria del Carmine and the parents of the newborns involved in the study for their collaboration and patience, Mariapia Giuseppa Piccininni and Roberta Eccher for help in collecting the data, and Sampsa Vanhatalo and Anton Tokariev for generously sharing their realistic newborn head model. This work was supported by grants from the European Research Council under the European Union’s Seventh Framework Programme (FP7/2007-2013)/Advanced Grant ERC PREMESOR G.A. [nr. 295517], from the Fondazione Caritro Grant Biomarker DSA, and from the PRIN 2015 (Neural bases of animacy detection, and their relevance to the typical and atypical development of the brain) to G.V.

